# The AFB1 auxin receptor controls the cytoplasmic auxin response pathway in *Arabidopsis thaliana*

**DOI:** 10.1101/2023.01.04.522696

**Authors:** Shiv Mani Dubey, Soeun Han, Nathan Stutzman, Michael J Prigge, Eva Medvecká, Matthieu Pierre Platre, Wolfgang Busch, Matyáš Fendrych, Mark Estelle

## Abstract

The phytohormone auxin triggers root growth inhibition within seconds via a non-transcriptional pathway. Among members of the TIR1/AFBs auxin receptor family, AFB1 has a primary role in this rapid response. However, the unique features that confer this specific function have not been identified. Here we show that the N-terminal region of AFB1, including the F-box domain and residues that contribute to auxin binding, are essential and sufficient for its specific role in the rapid response. Substitution of the N-terminal region of AFB1 with that of TIR1 disrupts its distinct cytoplasm-enriched localization and activity in rapid root growth inhibition. Importantly, the N-terminal region of AFB1 is indispensable for auxin-triggered calcium influx which is a prerequisite for rapid root growth inhibition. Furthermore, AFB1 negatively regulates lateral root formation and transcription of auxin-induced genes, suggesting that it plays an inhibitory role in canonical auxin signaling. These results suggest that AFB1 may buffer the transcriptional auxin response while it regulates rapid changes in cell growth that contribute to root gravitropism.

## Introduction

Auxin rapidly inhibits root growth via a non-transcriptional signaling pathway. This rapid growth response is critical for gravitropism and is accompanied by several cellular responses such as apoplastic alkalization, membrane depolarization and very rapid Ca^2+^ influx into the cytoplasm^1,2^. Among members of the TIR1/AFB auxin receptor family, AFB1 was reported to mediate the rapid root growth inhibition^3,4^. Further, the loss of AFB1 alone was sufficient to result in a significant defect in rapid root growth inhibition^3,4^, indicating its dominant role in this process.

## Results and Discussion

Consistent with these findings, we confirmed that the *afb1* mutant is resistant to auxin during the first 20 minutes of treatment whereas the *tir1* mutant is similar to wild type (Fig. 1a, b). As time progressed, the level of *afb1* mutant resistance decreased while that for *tir1* increased (Fig. 1b). This behavior is consistent with the roles of AFB1 and TIR1 in the nongenomic and transcriptional response, respectively. To test the role of receptors other than AFB1 in the rapid response, we measured auxin-induced root growth inhibition and the dynamics of the gravitropic response in the *tir1afb2* and *tir1afb345* mutants. Both the mutants responded to IAA similarly to wild type (Fig 1c). Previously, the *tir1afb345* mutant was shown to be resistant in the early phase^3^. The basis for this difference is unknown but it might be related to the highly variable phenotype of *tir1afb345* seedlings. Consistent with the growth inhibition, the *afb1* mutant showed a delay in early gravitropic response, however *tir1afb2* and *tir1afb345* mutants responded with a similar dynamics to the Col-0 control (Fig. 1d; Supp. movie 1).

**Figure 1:**
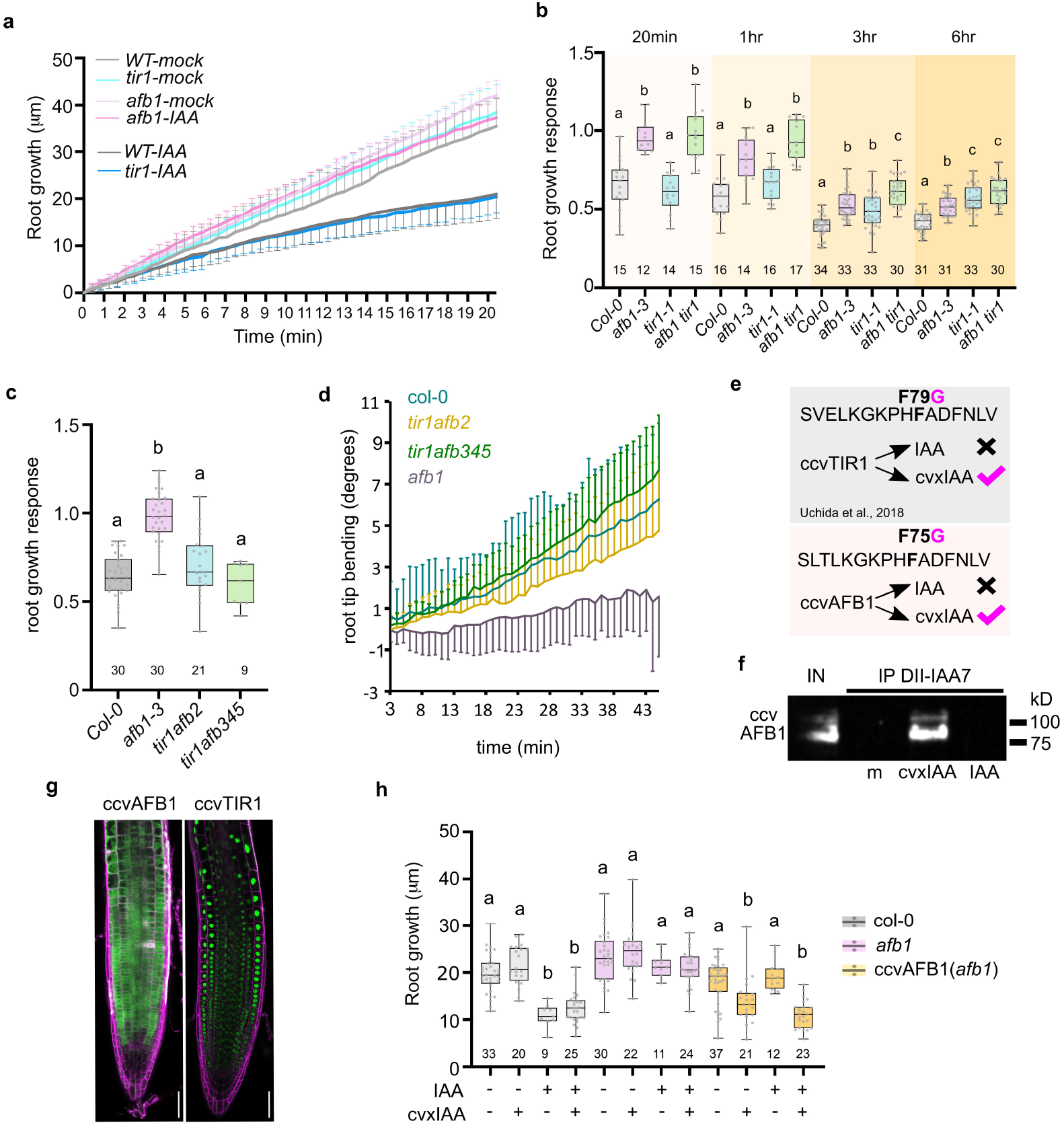
AFB1 triggers the early phase of auxin-dependent rapid root-growth inhibition. a) Root growth (μm) of Col-0, *afb1-3*, and *tir1-1* seedlings in response to mock (ethanol) or IAA treatment (10 nM). Root growth was measured every 25 seconds for 20 minutes. Col-0-mock (n=11), Col-0-IAA (n=13), *afb1-mock* (n=7), *afb1-IAA* (n=11), *tir1*-mock (n=12), *tir1*-IAA (n=12). Mean ± s.d. (s.d. represented as error bars). b) Root growth response (growth of treated roots normalized to growth of the respective mock-treated roots) to IAA (10 nM) for 20 minutes, 1 hr, 3hr and 6hr. c) Root growth response to IAA (10nM; 15 min) of Col-0, *afb1, tir1afb2*, and *tir1afb345* seedlings. d) Gravitropic response of Col-0, *afb1, tir1afb2*, and *tir1afb345* roots. Mean root tip angle ± s.d. (represented as error bars), time after a 90° gravistimulation is indicated, n=10-14 individual roots. e) Schematics of ccvAFB1 receptor design. Highlighted amino acid is the substitution in TIR1 and AFB1 that confers cvxIAA binding. f) In vitro pull-down assays of ccvAFB1-mScarlet by IAA7-DII peptide co-incubated with mock (m) 10 μM cvxIAA or 10 μM IAA. Input is shown (IN), ccvAFB1-mScarlet detected by anti-mCherry antibody. Estimated molecular weight of ccvAFB1-mScarlet = 92 kD. Uncropped membrane is shown in supplementary figure 4. g) Subcellular localization of *ccvAFB1-mScarlet* and *ccvTIR1-mScarlet* (green) in Arabidopsis root tips counterstained with propidium iodide (magenta); both constructs controlled by *pTIR1* promoter. Scale bars = 50 μm. h) Root growth after 15-minute treatment with mock, IAA (10 nM), cvxIAA (500 nM) or their combination in Col-0, afb1-3, and *ccvAFB1* (*afb1-3*) Only *ccvAFB1* roots respond to cvxIAA. Boxplots in b, c, h represent the median and the first and third quartiles, and the whiskers extend to minimum and maximum value; all data points are shown as dots. Statistical difference according to Ordinary one-way ANOVA coupled with Tukey’s multiple comparison tests (p<0.05) indicated by letters.

To exert greater control of the rapid auxin response, we prepared *Arabidopsis thaliana* lines expressing fluorescent protein-tagged versions of the synthetic receptor-ligand system – ccvTIR1 and ccvAFB1 controlled by the *pTIR1* promoter. The ccv (concave) receptor versions are ‘blind’ to the natural auxin IAA. Instead, they bind the synthetic cvxIAA (convex IAA^5^) (Fig. 1e,f), and show similar subcellular localization to the native proteins (Fig. 1g). The ccvAFB1 protein was sufficient to trigger the rapid root growth inhibition when seedlings were treated with cvxIAA (Fig. 1h). In previous studies it was shown that the cvxIAA-ccvTIR1 pair triggers a response that is several minutes delayed in comparison to the effect of natural IAA^2,6^. In our experiments, the cvxIAA – ccvTIR1 pair triggers a response that is significantly slower^4^ than the cvxIAA – ccvAFB1 system. Taken together, these results clarify previous discrepancies regarding the overlapping function of F-box receptors in the context of rapid auxin responses and suggest that AFB1 is the primary receptor for the rapid auxin responses.

One of the earliest detectable responses to auxin in the root is the influx of calcium^7^. A mutant lacking the CNGC14 calcium channel shows a delay in the gravitropic response^1^ that resembles the *afb1* gravitropic defect^4^, hinting at a role for AFB1 in auxin-dependent calcium influx. We therefore visualized cytosolic calcium levels in the *afb1* mutant using the R-GECO1 sensor^8^ and vertical microfluidic microscopy^4^. The control line showed an almost immediate elevation in cytosolic calcium (calcium transient) in response to 150 nM IAA, as described before^1,9,10^ (Fig. 2a, b). Strikingly, in *afb1* roots, cytosolic calcium did not increase after auxin application; instead, a mild calcium increase occurred only ca. 240 seconds after the treatment (Fig. 2a,b, Supp. movie 2). This indicates that the AFB1 receptor is required for the auxin-induced calcium transient.

**Figure 2:**
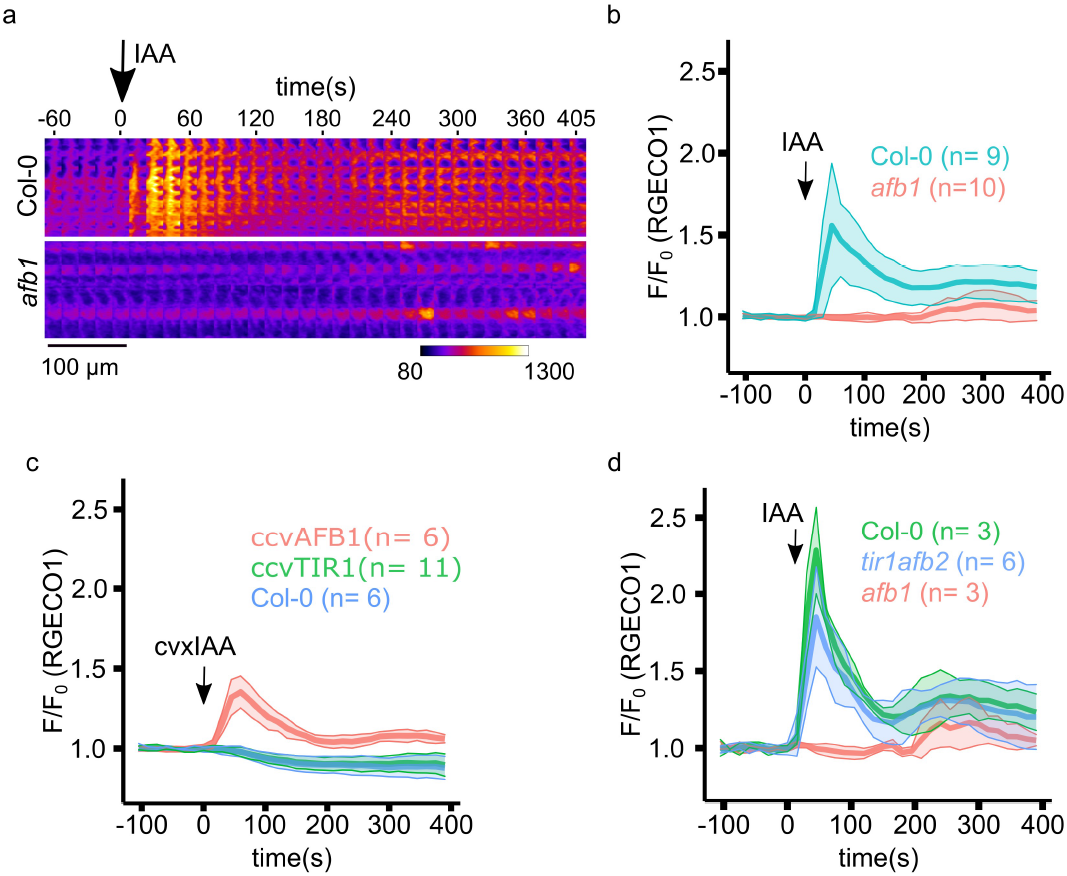
AFB1 controls auxin-dependent calcium transients in Arabidopsis primary roots. a) A kymograph showing auxin-induced R-GECO1 intensity increase indicating cytosolic calcium ([Ca^2+^]_cyt_) transients after application of 150 nM IAA (arrow) in Col-0 and *afb1-3* root epidermal cells. Fluorescence intensity look-up table is indicated. b-d) Quantification of R-GECO1 intensity indicating [Ca^2+^]cyt transients. b) Response to 150 nM IAA (arrow) in Col-0 and *afb1-3* root epidermal cells. c) Response of *ccvAFB1-mVenus, ccvTIR1-mScarlet* and Col-0 root epidermal cells to 500 nM cvxIAA. d) Response of *tir1afb2* root epidermal cells to 150 nM IAA, shown with positive (Col-0) and negative control (*afb1-3*). b-d) Normalized mean R-GECO1 fluorescence intensity F/F0 ± s.d. (represented as shaded areas). Auxin treatment is indicated by arrows.

To test whether AFB1 is upstream and sufficient for the calcium transient, we introduced the RGECO-1 sensor into the *ccvAFB1* and *ccvTIR1* lines. While in both the control and the *ccvTIR1* line, the application of 500 nM cvxIAA did not elicit a detectable calcium transient (Fig.2c), the compound triggered an immediate calcium transient in the *ccvAFB1* line (Fig.2c, Supp. movie 3). In contrast to these results, cvxIAA has been reported to trigger a calcium spike in a ccvTIR1 line expressing the GCaMP3 sensor^11^. We speculate that cvxIAA might trigger a transient detectable by GCAMP. However, the difference between control, ccvTIR1 and ccvAFB1 R-GECO-1 lines clearly shows that ccvAFB1 is required and sufficient for cytosolic calcium increase (Fig.2).

Finally, to determine if AFB1 is sufficient to trigger calcium influx in the case of the native receptor, we analyzed IAA-induced calcium transient in the *tir1afb345* and *tir1afb2* mutants. Unfortunately, the RGECO-1 construct was silenced in the *tir1afb345* line. On the other hand, the *tir1afb2* mutant responded to IAA treatment with a calcium transient comparable to the wild-type control (Fig. 2d, Supp. movie 4). These results show that AFB1 is upstream of the auxin-induced calcium transient that triggers the rapid growth responses including early root gravitropic responses^1,4,9^. It is intriguing that AFB1 has recently be shown to have adenyl cyclase activity^11^. Thus, it is possible that AFB1-mediated cAMP production triggers the activity of the CNGC14 channel.

Despite the apparent differences in their modes of action, TIR1 and AFB1 are the most recently diverged members of the TIR1/AFB family in *Arabidopsis^3^*. To determine if their functional specificity is related to their expression pattern, we expressed TIR1 in the *AFB1* expression domain. The *pAFB1:TIR1-mCitrine* expression pattern mimicked the AFB1 expression pattern^3^ (Fig. S1a). However, the transgene failed to rescue the *afb1* phenotype (Fig. S1b) even though the levels of AFB1 and TIR1 proteins were similar in these lines (Fig. S1g). As expected, the *pAFB1:AFB1-mCitrine* transgene *(gAFB1* #1 and *gAFB1 #2)* complemented the *afb1* phenotype. Interestingly, one of the *AFB1* complementation lines, *gAFB1 #2*, exhibited a higher expression level than either the wild-type or *gAFB1 #1* (Fig. S1c,d) and showed a hypersensitive rapid response to auxin (Fig. S1e). These data demonstrates that the functional differences between TIR1 and AFB1 are related to differences in their protein sequences, rather than expression pattern.

We previously showed that the TIR1/AFB proteins are partitioned between the cytoplasm and the nucleus. Interestingly, AFB1 is both the most abundant member of the family and highly enriched in the cytoplasm, while TIR1 is primarily nuclear^3^. To determine if subcellular localization is decisive for function in the rapid response, we added either a NUCLEAR LOCALIZATION SEQUENCE (NLS) or NUCLEAR EXCLUSION SEQUENCE (NES) to the AFB1-mCitrine receptor (*gAFB1-NLS* and *gAFB1-NES*). The resulting fusion proteins were highly enriched in the nucleus and cytoplasm respectively as expected (Fig. 3a). Intriguingly, only AFB1-NES rescued the phenotype of *afb1* (Fig. 3b), demonstrating that AFB1 must be localized to the cytoplasm to mediate rapid root growth inhibition. In a complementary approach, we attempted to generate *gTIR1-NES-mCitrine* plants but failed to recover lines with significant levels of TIR1 accumulation despite the presence of high transcript levels (Fig. S1f,g). It is possible that TIR1 is particularly unstable in the cytoplasm.

**Figure 3:**
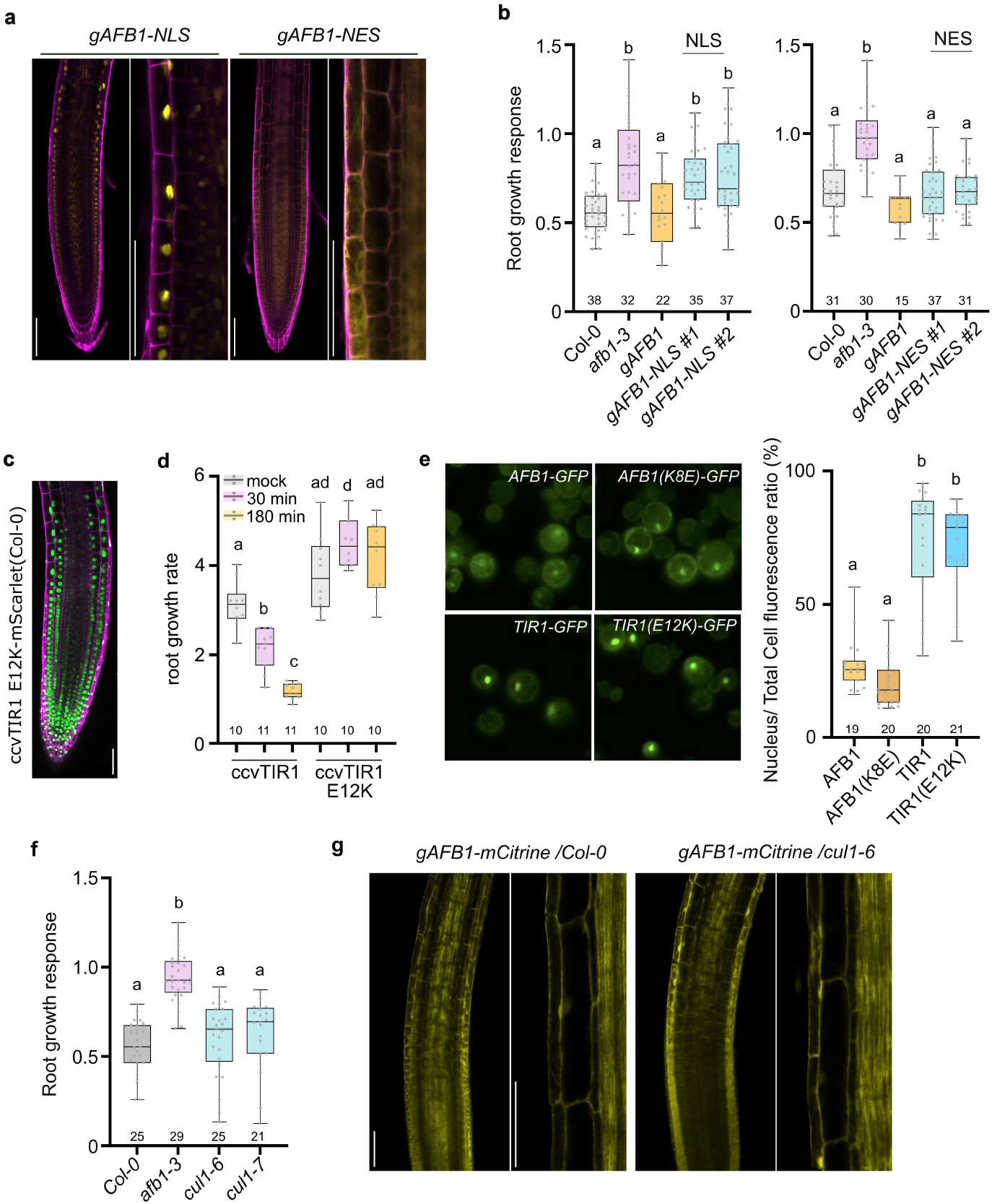
Cytoplasmic AFB1 regulates rapid root growth. a) Subcellular localization of AFB1-NLS-mCitrine and AFB1-NES-mCitrine (yellow) in *afb1-3;* roots stained with propidium iodide (magenta). Scale bars = 100 μm. b) Root growth response (growth of treated roots normalized to growth of the respective mock-treated roots) of Col-0, *afb1-3, gAFB1* (*afb1-3*) and *gAFB1-NLS/NES #1, #2* (*afb1-3*) to IAA (10 nM; 20 minutes). c) The ccvTIR1-E12K-mScarlet (Col-0) protein (green) localizes to nuclei. Stained with propidium iodide (magenta), scale bar= 50 μm. d) Root growth rate of *ccvTIR1-mScarlet* (Col-0) and *ccvTIR1-E12K-mScarlet* (Col-0) in mock or 500 nM cvxIAA treated seedlings. e) Subcellular localization of AFB1, AFB1(K8E), TIR1, and TIR1(E12K)-GFP in Col-0 *Arabidopsis* protoplasts (left) and quantification of relative nuclear localization (right), calculated as the ratio of nuclear fluorescence to total cell fluorescence. f) Root growth response of Col-0, *afb1-3, cul1-6* and *cul1-7* to IAA (10 nM; 20 minutes). g) Expression pattern and subcellular localization of AFB1-mCitrine in Col-0 and *cul1-6* background. Scale bar= 100 μm. Boxplots in b, d, e, f represent the median and the first and third quartiles, and the whiskers extend to minimum and maximum value; all data points are shown as dots. Letters indicate statistical differences according to Ordinary one-way ANOVA coupled with Tukey’s multiple comparison tests (p<0.05).

It has been reported that a polymorphism within the F-box domain of AFB1 (K at position 8 rather than E) strongly reduces its ability to interact with CUL1 and assemble into an SCF complex^12^. The TIR1 E12K mutation, recapitulating AFB1, significantly reduces the interaction with CUL1 and results in a strong auxin-resistant phenotype^12^. To test whether the differential affinity of TIR1 and AFB1 for CUL1 determines their distinct subcellular localization and function, we prepared the E12K version of ccvTIR1 to mimic the weak binding affinity of AFB1 with CUL1. Surprisingly, the protein still localized to the nucleus (Fig. 3c), interacted with the degron domain of Aux/IAA7 in a cvxIAA-dependent manner *in vitro* (Fig. S2b), but was unable to inhibit root growth in response to cvxIAA (Fig. 3d). We also expressed TIR1 E12K and AFB1 K8E in Arabidopsis protoplasts; these amino acid substitutions did not impact cellular localization (Fig.3e). These results indicate that the association with the SCF complex is not required for the co-receptor assembly and does not determine cellular localization.

As AFB1 does not bind CUL1 efficiently, we tested the possibility that AFB1 functions independently of an SCF complex by examining the rapid auxin response in the *cul1-6* and *cul1-7* mutants. *CUL1* is an essential gene during embryogenesis but these two hypomorphic alleles are viable and auxin resistant in a long-term root growth assay^13,14^. Both the *cul1-6* and *cul1-7* mutants exhibited a normal rapid response (Fig. 3f). In addition, AFB1 localization was not altered in the *cul1-6* mutant, confirming that AFB1 localization is not regulated by interaction with CUL1 (Fig. 3g). Similarly, recently published proteomic data^2^ confirms that TIR1, but not AFB1, interact with CUL1 in an IP-MS experiment, despite the relative abundance of AFB1. In contrast, both TIR1 and AFB1 interact with ASK1 as expected since this had been shown previously^12^. These results suggest that CUL1 binding and presumably SCF complex formation are not required for AFB1-triggered rapid root growth inhibition.

The well-known substrates for SCF^TIR1/AFB^ are the Aux/IAA transcriptional repressors. AFB1 has been shown to interact with several members of this family, either in Y2H assays or in plants^15^. We tested the *axr2-1* and *shy2-2* mutants in the rapid response assay. These two lines have mutations in the DII region of IAA7 and IAA3 respectively, that act to stabilize the protein and confer auxin resistance in long term root growth assays^16,17^. Neither mutant exhibited a significant change in rapid root growth inhibition (Fig. S1h), suggesting that Aux/IAA proteins do not contribute to the rapid response. Although these results suggest that the Aux/IAAs may not be involved in the rapid response, it is important to note that there are 28 members in the family and it is possible that one or more of these have specialized function in the cytoplasm.

To identify the domains responsible for AFB1-specific localization and function, we generated a set of *TIR1/AFB1* domain swap constructs under control of the *AFB1* promoter and introduced them into the *afb1* mutant. The chimeric proteins are named according to the origin of each segment; T for TIR1, and A for AFB1 (Fig. 4a). As shown earlier, AFB1 (or AAAA) was localized to both nucleus and cytoplasm, and rescued the *afb1* phenotype, while TIR1 (TTTT), was largely localized to the nucleus and failed to restore the *afb1* defect (Fig. 4b,c). Interestingly, among the 4 chimeric proteins, only TAAA was abundant in the nucleus similar to TIR1, and failed to restore the *afb1* mutant sensitivity to auxin. The ATAA, AATA, and AAAT chimeric proteins localized to both the nucleus and cytoplasm (Fig. 4b) and restored auxin sensitivity to the mutant (Fig. 4c). These results indicate that the N-terminal segment of AFB1 is important for AFB1’s cytoplasmic localization and function. This region includes the F-box domain that mediates the interaction between F-box proteins, ASK proteins and CUL1, and the N-terminal part of the Leucin-rich repeat domains (LRR) that participates in auxin and/or Aux/IAA binding^15,18^ (Fig. 4a). We therefore created two additional chimeric proteins. One contained the entire region 1 (iATTT) while the second contained only the F-box from AFB1 (fbATTT) (Fig. 4a) and expressed them under the control of *pAFB1* and *pTIR1* promoters, respectively. Note that both *pTIR1* and *pAFB1* are active in the epidermis^3^ and iATTT and fbATTT are both localized to nucleus and cytoplasm (Fig.4d). However, only iATTT rescued the *afb1* phenotype (Fig. 4e), and elicited a calcium transient similar to that of wild-type (Fig. 4f,g; Supp. movie 5). The iATTT showed a patchy expression, and interestingly, we only observed calcium transients in iATTT expressing cells demonstrating the cell-autonomous nature of the auxin-triggered calcium transient. To corroborate the results, we created the ccv-fbATTT version of the receptor, which also showed cytoplasmic and nuclear localization (Fig. S2a), and interacted with the degron domain of Aux/IAA7 in a cvxIAA-dependent manner *in vitro* (Fig. S2b). However, like fbATTT, ccv-fbATTT failed to trigger root growth inhibition in response to cvxIAA (Fig.S2c). Finally, consistent with the previous results, the fbATTT line failed to restore the early gravitropic response of *afb1* mutant, while iATTT almost completely recovered the response (Fig. 4g, Supp.movie6). The lack of full complementation can be explained by the patchy expression of iATTT in the root tip (Fig. 4f).

**Figure 4.**
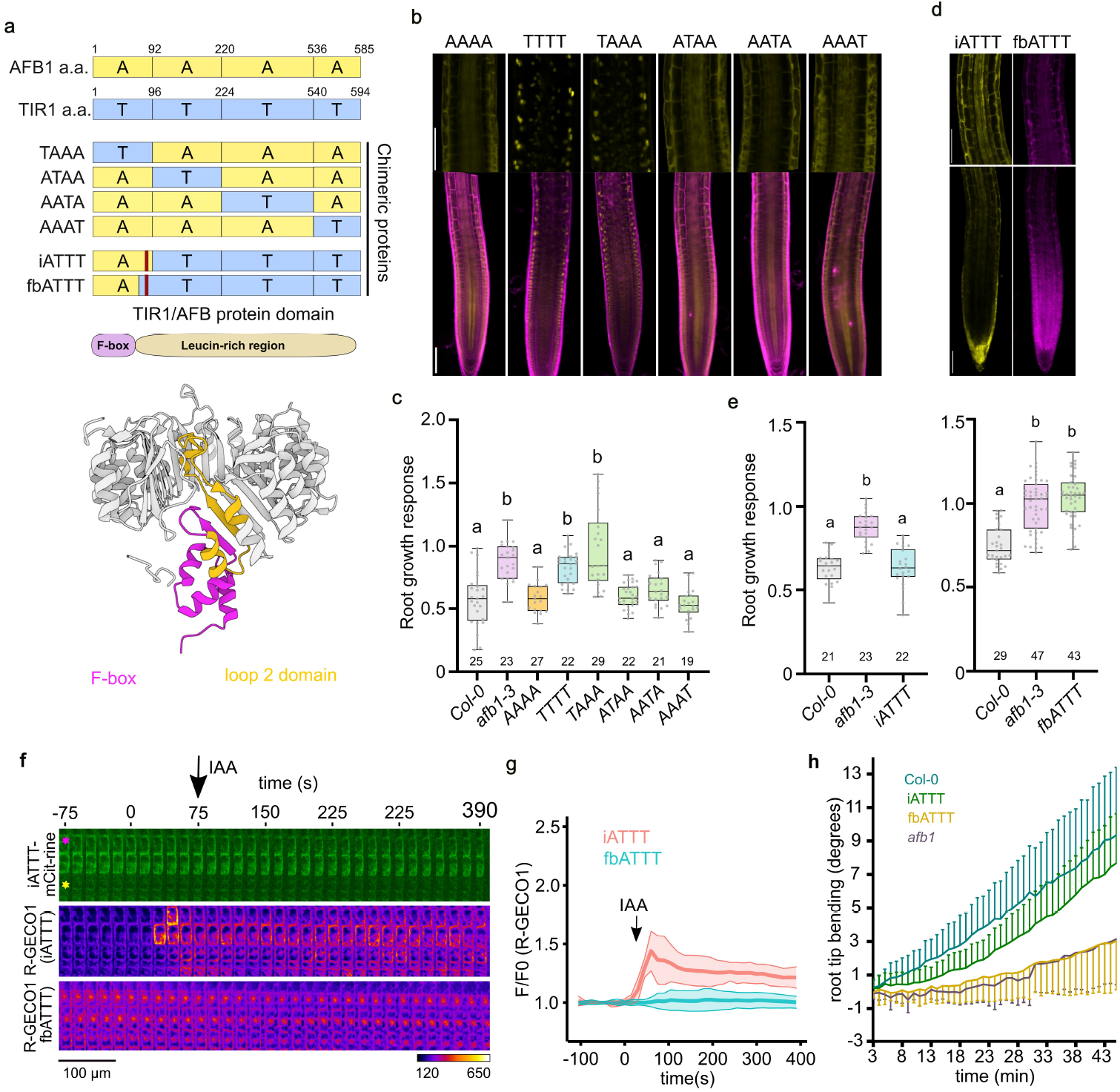
The N-terminal region of AFB1 is crucial for its role in the rapid auxin response. a) Schematic diagram of domain swap AFB1 (A-yellow) and TIR1 (T-blue) constructs. Numbers indicate amino acid position. Chimeric iATTT protein contains the F-box domain (magenta in 3D structure) and adjacent sequences in the LRR region (yellow in 3D structure) from AFB1, while the chimeric fbATTT protein contains only the AFB1 F-box domain (magenta in 3D). b) Expression pattern and subcellular localization of AFB1 (AAAA), TIR1 (TTTT) and chimeric proteins (TAAA, ATAA, AATA, AAAT). All domain swap proteins were regulated by the *AFB1* promoter in the *afb1-3* background. Scale bar=100 μm, stained with propidium iodide (magenta). c) Root growth response of Col-0, *afb1-3*, and domain swap lines to IAA (10 nM, 20 minutes); growth of treated roots normalized to growth of the respective mock-treated roots. d) Expression pattern and subcellular localization of iATTT-mCitrine and fbATTT-mScarlet proteins. The iATTT and fbATTT proteins were regulated by the *AFB1* and *TIR1* promoters respectively in the *afb1-3* background. Scale bar=100 μm. e) Root growth response of Col-0, *afb1-3, iATTT(afb1-3)* and *fbATTT(afb1-3)* roots to IAA (10 nM) for 20 minutes. f) A kymograph showing auxin-induced cytosolic calcium transients (R-GECO1 intensity) during application of 150 nM IAA (arrow) in iATTT(*afb1*) and fbATTT(*afb1*) root epidermal cells. On the top, the iATTT-mCitrine channel highlights the patchy expression of the construct, note that the iATTT-expressing cell (asterisk) also shows high calcium transient. Fluorescence intensity look-up table is indicated. g) Quantification of R-GECO1 intensity indicating [Ca^2+^] cyt transients in iATTT and fbATTT root epidermal cells. IAA treatment (150 nM) shown by an arrow. n = 6-11 Boxplots in c,e represent the median and the first and third quartiles, and the whiskers extend to minimum and maximum value; all data points are shown as dots. h) Quantification of the gravitropic response in Col-0 (n=20), *afb1* (n=12), *iATTT* (n=23), and *fbATTT* (n=23) roots. Mean root tip angle ± s.d. (represented as error bars) is shown. Time after 90° gravistimulation is indicated. Boxplots in c, e represent the median and the first and third quartiles, and the whiskers extend to minimum and maximum value; all data points are shown as dots. Letters indicate statistically different groups according to Ordinary one-way ANOVA coupled with Tukey’s multiple comparison tests (p<0.05).

These results show that the F-box domain determines the cytoplasmic/nuclear partitioning of the receptors; that the cytoplasmic localization of the auxin receptor is required but not sufficient for function in the rapid response; and, finally, that the sequences in the N-terminal part of the AFB1 LRR domain are required for the function in the rapid auxin response. Since a clear nuclear-localization signals (NLS) is not present in the F-box domain of TIR1, it is possible that an unknown protein interacting with the F-box domain is involved in the regulation of subcellular localization.

Since AFB1 is abundant in both the cytoplasm and nucleus, we also determined the effects of manipulating AFB1 levels on long term auxin responses that are mediated by canonical auxin signaling. In a long-term root growth assay, we found that the *gAFB1-NLS* line displayed significant auxin resistance, while the *gAFB1-NES* line was slightly hypersensitive (Fig. 5a, S3a). Auxin-hypersensitivity of the line expressing AFB1-NES suggests that the rapid AFB1-dependent pathway can also affect auxin response over a longer time frame. However, we could not exclude the possibility that cytoplasmic AFB1 also negatively affects canonical auxin signaling because this effect could be masked by the AFB1-mediated root growth inhibition.

**Figure 5.**
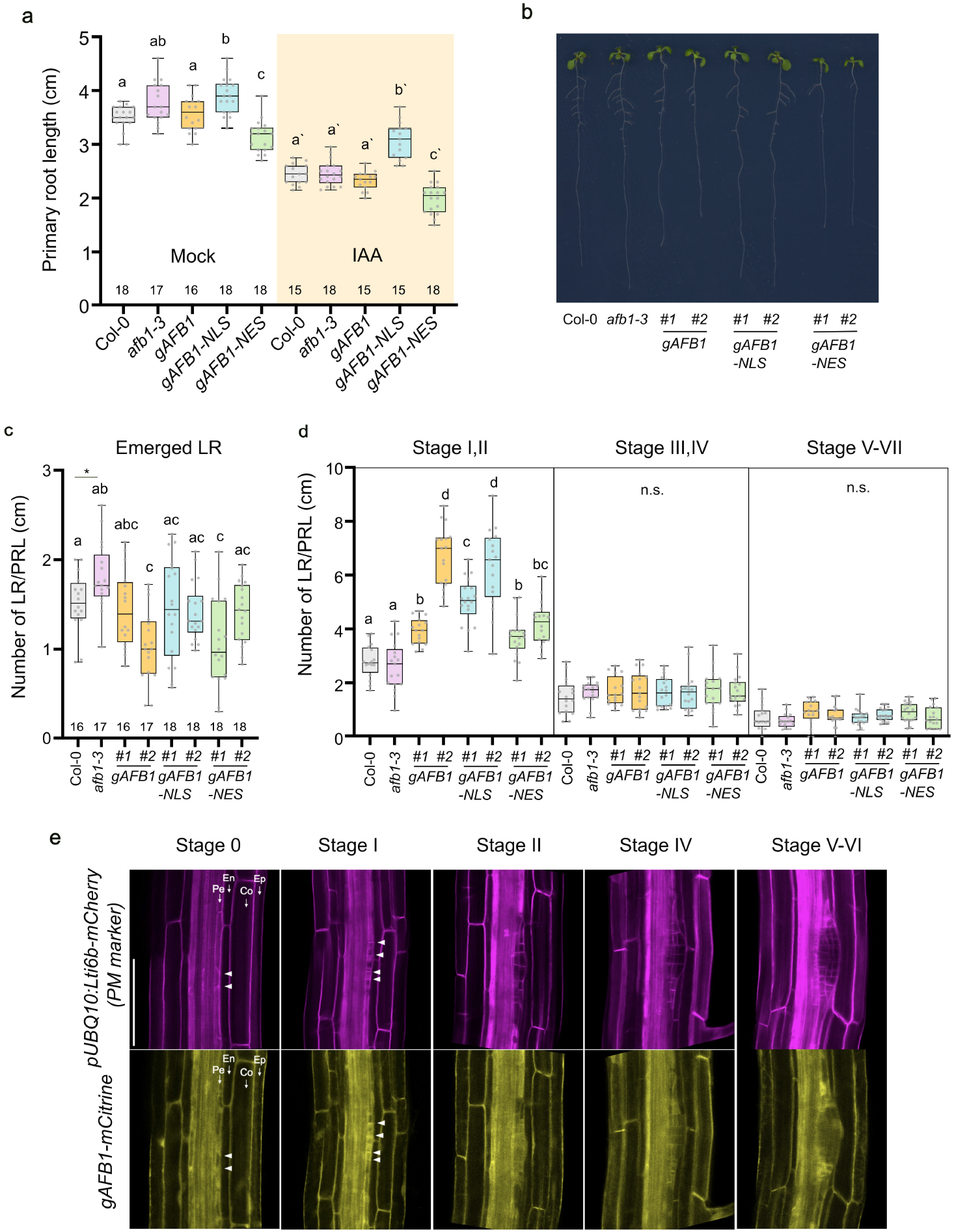
AFB1 negatively regulates canonical auxin signaling. a) Primary root length of five-day-old seedlings of Col-0, *afb1-3, gAFB1* (*afb1-3*), and *gAFB1-NLS/NES* (*afb1-3*) lines treated with either 100 nM IAA or mock (ethanol) for 3 days. b) Lateral root phenotype in nine-day-old seedlings of Col-0, *afb1-3, gAFB1* (*afb1-3*), and *gAFB1-NLS/NES* (*afb1-3*) lines. c) Number of emerged lateral roots per primary root length in Col-0, *afb1-3, gAFB1* (*afb1-3*) and *gAFB1-NLS/NES* (*afb1-3*) lines (#1 and #2 indicates two independent lines). d) Number of non-emerged primordia at different stages in lateral root developed expressed per primary root length in Col-0, *afb1-3, gAFB1 (afb1-3)* and *gAFB1-NLS/NES* (*afb1-3*) lines (#1 and #2 indicates two independent lines). e) Expression pattern of gAFB1-mCitrine; pUBQ10:Lti6b-mCherry during lateral root development. pUBQ10:Lti6b-mCherry was used as a plasma membrane marker. Ep; Epidermis, Co; Cortex, En; Endodermis, Pe; Pericycle. Scale bar=100 μm. Box plots represent the median and the first and third quartiles, and the whiskers extend to minimum and maximum value; all data points are shown as dots. Letters above box plots indicate statistical differences according to Ordinary one-way ANOVA coupled with Tukey’s multiple comparison tests (p<0.05).

Earlier genetic studies suggested that AFB1 may be a negative regulator of LR formation^3^. Here we show that the *afb1* mutant produces slightly more lateral roots than wild type while two AFB1 complementation lines *(gAFB1 #1* and *#2*) produce many fewer lateral roots, confirming this hypothesis (Fig. 5b,c). Interestingly, the lines expressing AFB1-NLS and AFB1-NES also have reduced numbers of LR (Fig. 5b,c). All three lines, *gAFB1, gAFB1-NLS* and *gAFB1-NES*, exhibited significantly increased numbers of early stage LR (stages I-II) (Fig. 5d) compared to wild type indicating that AFB1 does not affect LR initiation, but rather suppresses LR emergence, when it is present in either the cytoplasm or nucleus. In addition, we observed that all three AFB1 proteins, AFB1, AFB1-NLS and AFB1-NES are broadly expressed in both developing primordia and its overlaying tissues (Fig. 5e, S3a).

To determine if the role of AFB1 during lateral root development is associated with auxin regulated transcription we examined the expression of the auxin-responsive genes *IAA5, IAA6, IAA19* and two lateral root-related genes, *LBD16* and *LBD29^19^*. As expected, auxin treatment increased expression of these genes in wild-type seedlings (Fig. S3b). Intriguingly, induction of *IAA5, IAA6* and *LBD29* was greater in *AFB1*, but suppressed in a dose-dependent fashion in the *AFB1* complementation lines (*gAFB1 #1* and *gAFB1 #2*). Moreover, this suppression was also observed in *gAFB1-NLS* (Fig. S3b). These results are consistent with the lateral root phenotype of these lines. Surprisingly, auxin induction of gene expression was also suppressed in *gAFB1-NES* seedlings. These data indicate that both nuclear and cytoplasmic AFB1 function as a negative regulator of auxin-mediated transcription, presumably leading to the inhibition of canonical auxin signaling during long-term development.

Taken together, we propose that while cytoplasmic AFB1 induces non-genomic rapid auxin response which is dependent on CNGC14-mediated Ca^2+^ signaling, both nuclear and cytoplasmic AFB1 inhibit canonical auxin signaling. In the case of nuclear AFB1, the protein may act as a dominant-negative in a manner similar to TIR1(E12K)^12^. In contrast, how cytoplasmic AFB1 acts to suppress the canonical pathway is unknown. Regardless, this activity may serve to integrate the two auxin responses as the root responds to changing environmental conditions.

## Acknowledgements

This work was supported by the National Institute of General Medical Sciences (NIGMS) with grants to ME (R35GM141892) and to WB (R01GM127759), and by the European Research Council (grant no. 803048) to MF. MPP was supported by a long-term postdoctoral fellowship (LT000340/2019 L) by the Human Frontier Science Program Organization. The authors thank Melanie Krebs and Karin Schumacher for providing the R-GECO1 plasmid, Nelson BC Serre for experimental guidance.

## Supplementary Figures

**Supplementary figure 1:**
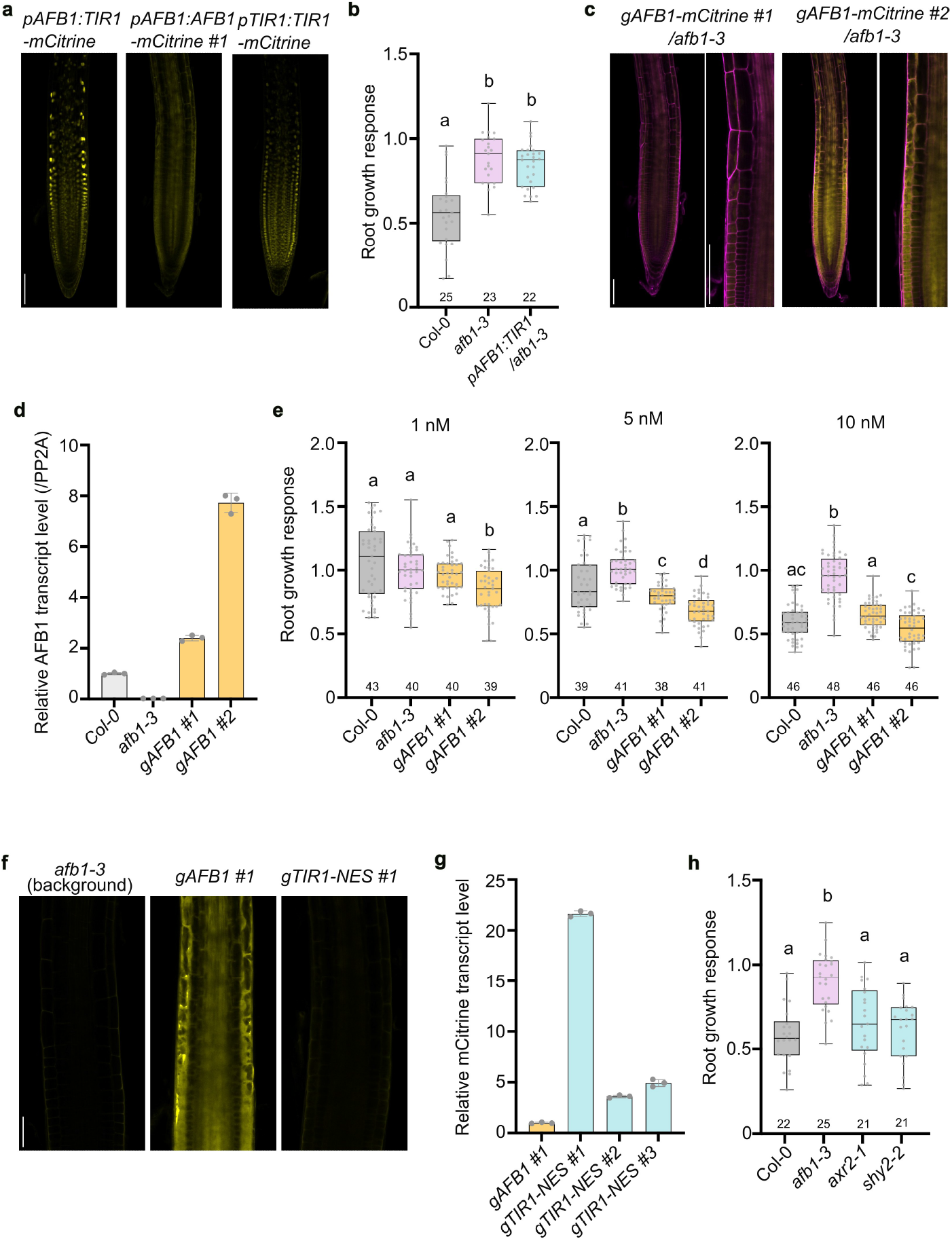
*afb1* phenotype is not rescued by TIR1 but AFB1 in dose-dependent manner. a. Expression pattern of *pAFB1:TIR1-mCitrine* (*afb1-3*), *pAFB1:AFB1-mCitrine* (*afb1-3*) and *ptir1:TIR1-mCitrine* (*tir1-1*) in the root. Scale bar=100 μm. b. Root growth response of Col-0, *afb1-3*, and *pAFB1:TIR1(afb1-3)* lines to IAA (10 nM; 20 minutes). c. d. Expression pattern of *gAFB1-mCitrine #1 and #2* (*afb1-3*) in the root (c) and the relative *AFB1* transcript level by qRT-PCR analysis (d). Scale bar=100 μm. Error bar indicates ±S.E.M. The cell walls were stained with propidium iodide (magenta). e. Root growth response of Col-0, *afb1-3, gAFB1 #1* and *#2* (*afb1-3*) to IAA (1 nM, 5 nM and 10 nM; 20 minutes). f, g. Expression of *gTIR1-NES-mCitrine* (f) and transcript levels (g) in *gTIR1-NES-mCitrine* (*afb1-3*) lines compared to *gAFB1 #1* (*afb1-3*). Fluorescent signal in f was over-exposed to detect very low fluorescence intensities. Error bar indicates ±S.E.M. Scale bar=100 μm. h. Root growth response of Col-0, *afb1-3, axr2-1* and *shy2-2* lines to IAA (10 nM; 20 minutes). In b, e, h, boxplots represent the median and the first and third quartiles, and the whiskers extend to minimum and maximum value; all data points are shown as dots. Letters indicate statistical differences according to Ordinary one-way ANOVA coupled with Tukey’s multiple comparison tests (p<0.05).

**Supplementary figure 2:**
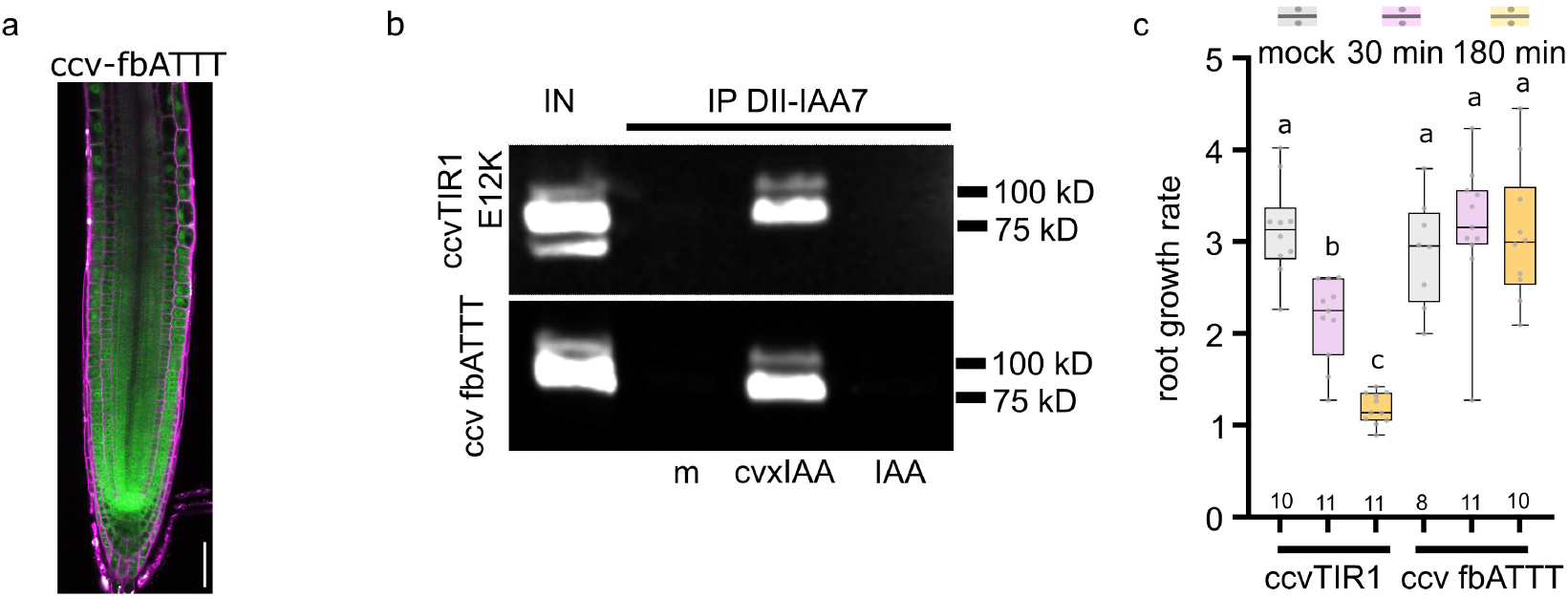
Cytoplasmic localization of ccv-fbATTT-mScarlet, and binding to cvxIAAis not sufficient to trigger root growth inhibition a. Subcellular localization of ccv-fbATTT-mScarlet (green) in Arabidopsis root tip counterstained with propidium iodide (magenta). Expression is under the control of pTIR1 promoter. Scale bar = 50μm. b. In vitro pull-down assay of ccvTIR1 E12K-mScarlet and ccv-fbATTT-mScarlet by IAA7-DII peptide co-incubated with mock (m) or 10 μM cvxIAA or 10 μM IAA. Input is shown (IN), ccvTIR1 E12K-mScarlet and ccv-fbATTT-mScarlet was detected by anti-mCherry antibody. Estimated molecular weights of both proteins are 92 kD. Number of replicates = 2. c. Root growth response of *ccvTIR1-mScarlet* (positive control) and *ccv-fbATTT-mScarlet* in mock or 500nM cvxIAA during 30 min and 180 min. Significantly different (p<0.05) statistical groups are indicated by letters, according to non-parametric ANOVA with Tukey’s multiple comparison.

**Supplementary figure 3:**
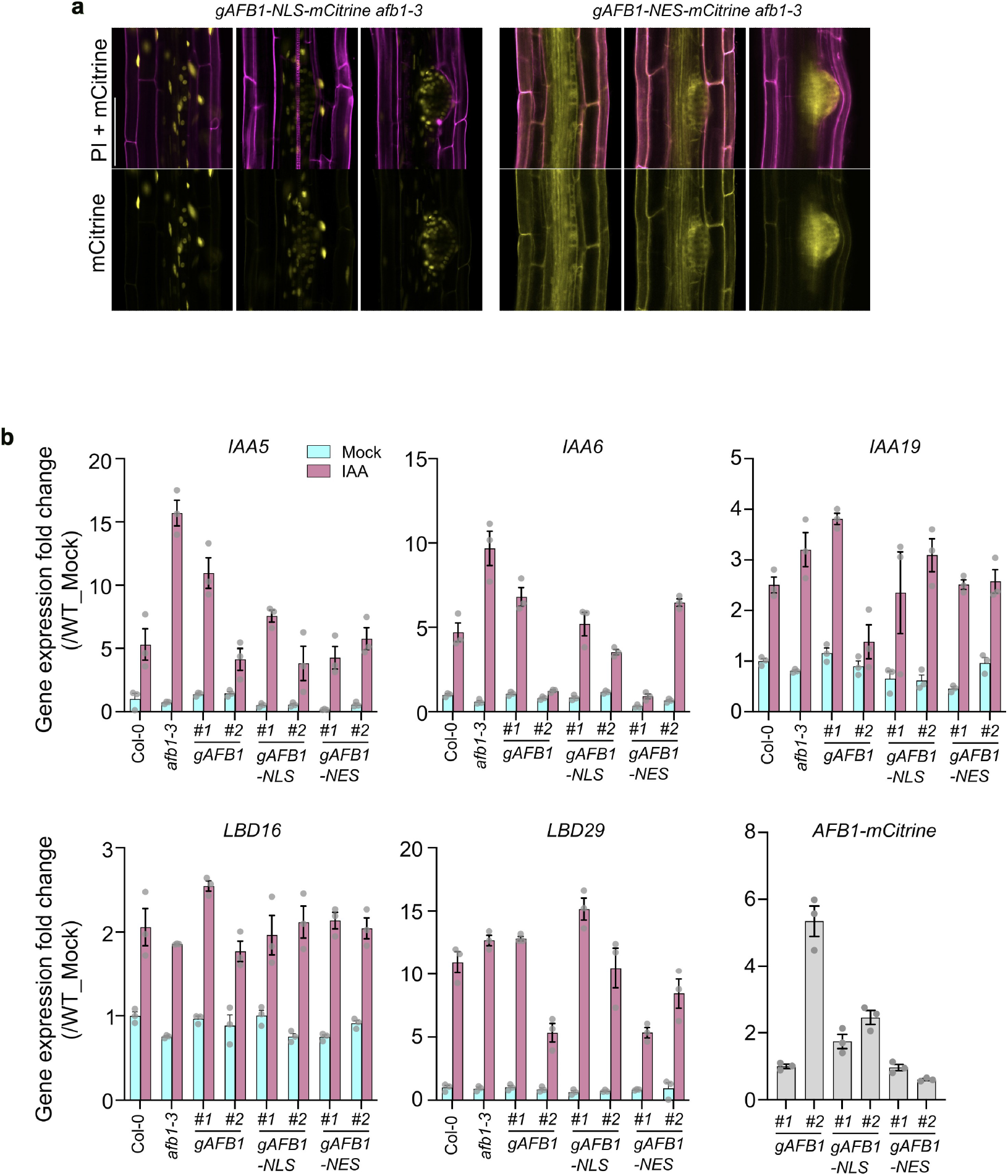
Both nuclear and cytoplasmic AFB1 negatively affect lateral root development and expression of auxin-responsive genes. a. Expression pattern of *gAFB1-NLS-mCitrine* (*afb1-3*) and *gAFB1-NES-mCitrine* (*afb1-3*) during lateral root development. Propidium Iodide (PI; magenta) was used as a cell wall marker. Scale bar = 100 μm. b. Relative expression of auxin-responsive genes (*IAA5, IAA6, IAA19, LBD16, LBD29*) in Col-0, *afb1-3, gAFB1 afb1-3, gAFB1-NLS afb1-3* and *gAFB1-NES afb1-3* lines in response to mock (ethanol) or IAA (100 nM) for 2 hr (#1 and #2 indicates two independent lines). The *mCitrine* transcript level was shown to compare *AFB1* transgene expression level between transgenic lines. Error bars indicate ±S.E.M.

**Supplementary figure 4.**
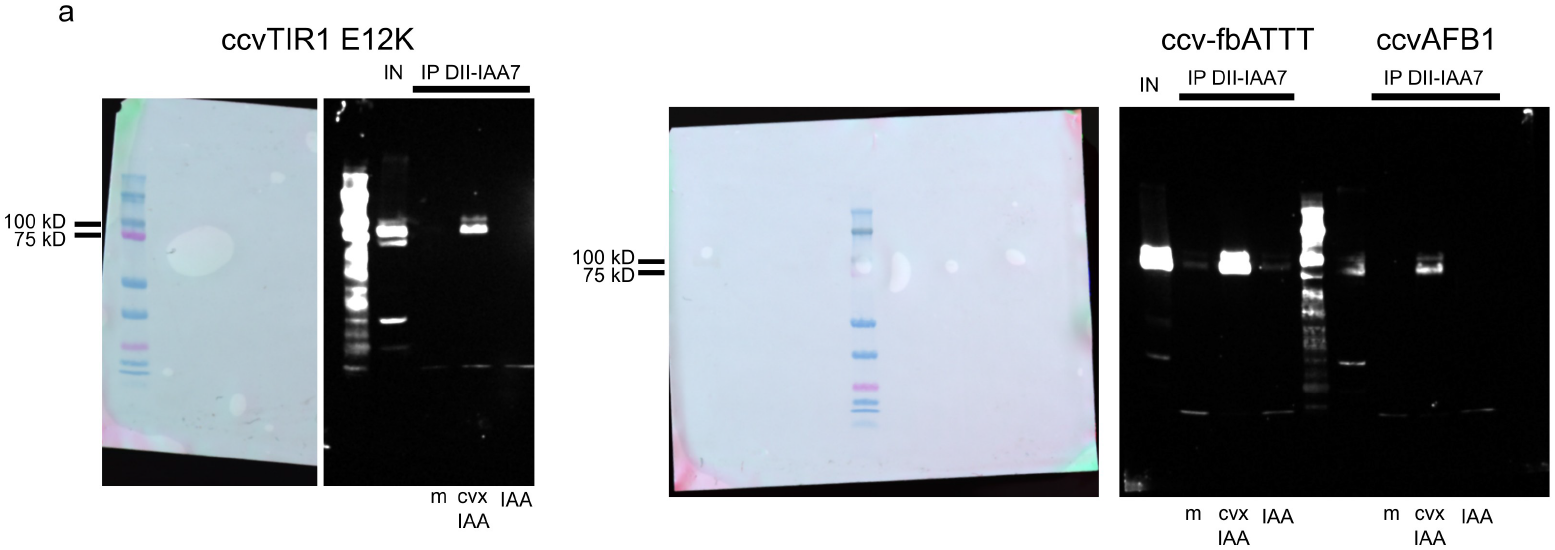
a. Full image of immunoblotted nitrocellulose membrane of images shown in Fig. 1f, Fig S2b. Brightfield images of membranes showing the prestained protein ladder are shown (on left). Mock (m), cvxIAA, IAA treatments are indicated below the respective lanes.

## Materials and Methods

### Plant Materials and growth conditions

All *Arabidopsis* mutants in this study were Col-0 background. The *afb1-3, tir1-1, tir1-1 afb1-3, tir1afb2, tir1 afb345* and *gAFB1-mCitrine* (*afb1-3*) were described and used previously^1^. The *shy2-2, axr2-1, cul1-6* and *cul1-7* were previously described^2–5^. Transgenic lines, *TIR1:ccvTIR1-mScarlet* (Col-0), *TIR1:ccvAFB1-mScarlet (afb1-3), PIN2:ccvAFB1-mVenus* (Col-0), *TIR1:fb-ATTT-mScarlet* (*afb1-3*), *TIR1:ccvfb-ATTT-mScarlet* (*afb1-3*), *TIR1:fb-ccvTAAA-mScarlet* (*afb1-3*), *TIR1:ccvTIR1 E12K-mScarlet* (Col-0) were generated using the GoldenBraid cloning system (see Molecular cloning). The R-GECO1 calcium sensor was introduced by crossing into *afb1-3, TIR1:ccvTIR1-mScarlet* (Col-0), *PIN2:ccvAFB1-mVenus* (Col-0), *TIR1:fb-ATTT-mScarlet* (*afb1-3*), and *AFB1:iATTT-mCitrine* (*afb1-3*). The *R-GECO1* (*tir1afb2*) and *R-GECO1* (*tir1afb345*) lines were generated by transformation of the respective mutants.

Seeds were sterilized with bleach (3% NaClO) and stratified in the dark cold room for 2-4 days. Plants were vertically grown on ½ MS media containing 0.8-1.0 % agar, 1% (w/v) sucrose, adjusted to pH 5.8 with KOH, and kept at 22 °C in a long day (16L/8D), growth room maintaining 60% humidity, and a light intensity of 100 μmol photons m^-2^ s^-1^.

### Molecular cloning

The *pMiniT-gAFB1(AT4G03190)-mCitrine* constructs were previously described^1^. The domain swap constructs were assembled using the NEBuilder HiFi DNA assembly kit (NEB). Briefly, *TIR1(AT3G62980)* domains or the entire *TIR1* 5’ UTR plus coding sequence were amplified with primers with 5’ extensions matching *AFB1*, then inserted into corresponding PCR-amplified *pMiniT-gAFB1* backbone fragments. These chimeric genes were then moved to the *pMP535* binary vector as *MluI-AscI* fragments. For generating *gAFB1-NES-mCitrine* and *gTIR1-NES-mCitrine* lines, a double-stranded oligo with the NES fragment was inserted into NheI-digested *pMiniT-gAFB1nhe* and *pMiniT-gTIR1NHE* constructs, in which the stop codons were mutated to create *NheI* restriction enzyme sites^1^. An *mCitrine* fragment was amplified, digested with XbaI, and then inserted into the *NheI* site after the *NES* fragment. For *pMiniT-gAFB1-NLS-mCitrine*, a fragment containing the *SV40 NLS* followed by *mCitrine* was amplified, cut with XbaI, and ligated into the NheI-digested *pMiniT-gAFB1nhe* plasmid. These *pMiniT-gAFB1/gTIR1-NES/NLS-mCitrine* were subcloned into the *pMP535* binary vector as described previously.

The first 53 and 57 amino acids of AFB1 and TIR1, respectively, ending in CYAVS were exchanged to generate the constructs fbATTT and fbTAAA, respectively. The ccvTIR1 and ccvfb-TAAA lines were generated by the substitution mutations F79 > G in TIR1^6^; ccvAFB1 and ccvfb-ATTT lines were generated by a substitution mutation F75 > G in AFB1. The ccvTIR1 E12K mutation was created as described before^7^. The plasmid constructs of *ccvTIR1, ccvAFB1, ccvfb-ATTT, ccvfb-TAAA, fbATTT, fbTAAA*, and *ccvTIR1 E12K* were first cloned into the pUPD2 vector. Then, the genes were cloned into *pDGB1alpha1* by combining the *TIR1* promoter, a C-terminal *mScarlet-I^8^* or *mVenus* tag, and a 35s terminator into the *pDGB1alpha1* vector. These transcriptional units were then combined with a Basta resistance cassette into the *pDGB3omega1* binary vector. Cloning steps were performed using the GoldenBraid method^9^ (https://gbcloning.upv.es/).

The binary constructs were transformed into *Agrobacterium tumefaciens* strain pGV3101 by electroporation, and then transformed into Col-0, *afb1-3, tir1afb2*, or *tir1afb345* plants by floral dipping^10^. The transgenic lines used in the study are summarized in Supplementary Table 1; all primers used for cloning are listed in Supplementary Table 2.

### Visualization of calcium transients

Calcium transients were visualized using the R-GECO1 reporter^11^ and imaged using a microfluidic setup combined with a vertical spinning disk microscope^12^. Five-day-old *Arabidopsis* seedlings were transferred to a sealable single-layer PDMS silicone chip. The PDMS silicone chip containing the seedlings was then placed on a vertical spinning disk microscope for 20 min to recover. The seedlings were imaged with a 20x/0.8 objective every 15 seconds for the first 5 minutes with control medium and then switched to treatment medium for 10 minutes. Constant media flow and switching between control and treatment medium was performed using OBI1 Elveflow software. Images were acquired with VisiView (Visitron Systems, v.4.4.0.14). Control and treatment medium were prepared with 1% (w/v) sucrose containing either 0 or 150 nM IAA. For *ccvAFB1* and *ccvTIR1 R-GECO1* lines, we used 500 nM cvxIAA (#M3141, BDL) prepared from a 10 mM stock in DMSO.

R-GECO1 time series images were analyzed in Fiji^13^. Images were stabilized by selecting a ROI in-root transition zone and using imageJ plugin registration followed by correcting 3D drift^14^. We then quantified the R-GECO1 intensity in a region of interest (ROI) in several epidermal cells in the root transition zone. Signal intensity at each timepoint (F) was normalized using the average value of 8 timepoints (2 min) before treatment (F0), obtaining the F/F0. In the case of *tir1afb2* mutant, the R-GECO1 expression was generally lower, therefore we subtracted the background noise level before calculating the F/F0.

### Quantification of gravitropic response

The gravitropic experiment was done as described^15^: a thin layer of growth medium (½ MS, 1% (w/v) sucrose in 1% plant agar, Duchefa) containing 5-day-old seedlings was placed into a 3D-printed microscopy chamber. The chamber was placed vertically on the vertical stage microscope for 45 minutes to recover. Then the chamber was rotated by 90 degrees. After the two minutes needed to set the imaging, the roots were imaged every minute for 43 minutes. The root tip angles were measured using ACORBA v1.2.

### Root growth assay

For the rapid root growth inhibition assay, we used two methods. The first method was previously described^1,16^, and was used for Figure 1a, 1b, 3b, 3f, 4c, 4e, S1b, S1e, S1h. Briefly, 10-15 five-day-old seedlings were transferred to culture chambers containing the growth medium supplemented with EtOH (mock) or 10 nM IAA (stock solution 10 μM in EtOH), and immediately imaged every 25 seconds for 20 minutes or every 72 seconds for 1 hr. The 50 images taken for 20 min or 1 hr were combined and processed by MatLab software using a customized Matlab script called Rootwalker^16^. Rootwalker MATLAB generated growth curves of individual roots (root length increment/time interval) for each treatment (mock and IAA) and exported to jpeg and excel files. In the second method, used in Figure 1c, h, five-day-old seedlings were transferred to growth medium containing either 0 or 10 nM IAA. The seedlings with the media were then immediately transferred to a 3D-printed microscopy chamber (24 × 60 mm) and placed on the vertical microscope. 4-5 min were required to set the stage positions on the microscope, and then roots were imaged for 10 minutes after each 5 min using a 5x/0.16 objective. Root growth (length increment) measured after stabilizing the background drift of the root tip using the Fiji plugin correct 3D drift, as described^12^.

For measuring long-term root growth (3 hr and 6 hr), vertically grown five-day-old seedlings were imaged at 1,200 dpi using a flatbed scanner at time 0, 3hr and 6hr after transferring onto treatment medium (mock and 10 nM IAA). Root tips of individual seedlings were marked at each time point, and root growth increment was measured using the segmented line tool in FIJI.

The root growth response to auxin was calculated as the length increment in treatment divided by the length increment in control conditions. The response value 1 indicates that the treatment did not affect root growth.

For measuring primary root length after long-term auxin treatment, five day-old seedlings were transferred to media containing the ethanol (mock) or 100 nM IAA and incubated for 3 days. The response to IAA was expressed as IAA-treated primary root length divided by the mean value of mock-treated primary root length.

### Pull down and western blotting

Seven-day-old seedlings were used for total protein extraction using extraction buffer (150 mM NaCl,100 mM Tris, 0.5% NP-40, pH 7.5), 1X concentration protease inhibitors (#11836170001, Merck), and 1mM PMSF (R 63672, P-lab). Pulldowns were done as described^17^ with the following modifications: The biotinylated Aux/IAA7 DII peptide (AKAQVVGWPPVRNYRKN, N-terminal biotin) was bound to streptavidin-agarose beads (Merck, #16-126). 250 μl of total protein extract was combined with 45 μl of DII-beads and either 0 μM or 10 μM cvxIAA or 10 μM IAA. The reaction was kept at 4 °C for 1 h at constant mixing and then washed 3 times with extraction buffer. The pull-down samples were then subjected to Western blotting. Rabbit Polyclonal Anti-mCherry (#ab167453, Abcam), and Goat anti-Rabbit IgG (H+L) Secondary Antibody (Invitrogen) were used for immunodetection. Detection of secondary antibody was performed by ECL (#AS18 SecondaryECL, Agrisera) staining followed by imaging with Azure 600 imaging system. Number of experimental replicates-ccvAFB1-mScarlet = 02, ccv-fbATTT-mScarlet = 02, ccvTIR1 E12K-mScarlet = 02

### Microscopy imaging

For imaging fluorescence signal in the primary root and lateral root, four or five-day-old seedlings were stained with a 10 μg/ml aqueous solution of propidium iodide (PI) for 1-2 minutes, washed with water, and mounted on a glass slide for imaging. PI (excitation, 561nm; emission, 642nm), mCitrine (excitation, 514nm; emission, 556nm) and mCherry (excitation, 594nm; emission, 628nm) signal were detected using a Plan-Apochromat 20x/0.8 NA air or 40x/1.2 NA WI objective lens on a Zeiss LSM 880. For imaging fluorescence signal in *Arabidopsis* protoplast, *AFB1-GFP, AFB1(K8E)-GFP*, *TIR1-GFP*, and *TIR1(E12K)-GFP* constructs under *CaMV35S promoter* were transfected into Col-0 *Arabidopsis* as previously described^18,19^. Transfected protoplasts were incubated in 12 well culture plates for 12 hr at room temperature and then imaged on glass slides using 20X/0.8 NA air objective lens on Zeiss 880 (GFP excitation, 488nm; emission, 507nm). Relative nuclear localization of AFB1, AFB1(K8E), TIR1, TIR1(E12K) in *Arabidopsis* protoplasts was calculated as the value of nuclear fluorescence signal divided by whole cell fluorescence signal in each protoplast cell (n=19-20). The fluorescence signal in the nucleus and whole cell was determined by Area*Mean grey value from each ROI to mark nucleus and whole cell region in FIJI software.

To acquire subcellular localization of ccvAFB1and ccvTIR1 E12K and ccv-fbATTT five-days old seedlings roots were stained with 2.5 μg/ml propidium iodide for 2 min and mounted on a glass slide with ½ MS, 1% sucrose medium followed by imaging with Zeiss LSM 880.

### Lateral root and primary root measurement

Vertically grown eight-day-old seedlings were used for lateral root measurements as previously described^20^. Briefly, seedlings were soaked in pre-heated 0.24 N HCl in 20% methanol, and further incubated at 60 °C for 10 minutes. The seedlings were then transferred to 7% NaOH, 7%hydroxylamine-HCl in 60% ethanol for 15 minutes at RT. The solution was replaced and incubated with a series of 40%, 20% and 10% ethanol for 5 minutes, and stored in 5% ethanol/ 25% glycerol solution. The root samples were mounted in 50% glycerol on glass slides and observed in bright field using a 40x/ 1.2 NA WI objective with a DIC filter on a Zeiss LSM 880.

### Gene expression analysis

Five-day-old seedlings vertically grown on ½ MS media were transferred to agar media containing ethanol (mock) or 100 nM IAA and further incubated in the growth chamber for 2 hr. Seedling pools (about 30 seedlings per genotype) from mock and IAA-containing media were harvested and immediately frozen in liquid nitrogen. Frozen seedling samples were used for RNA isolation using Qiagen RNeasy Plant Mini kit. 2 μg of RNA was used for RT-PCR using Superscript III RT-kit or Thermo Maxima H–master mix. qRT-PCR was performed using Bio-Rad SsoAdvanced SYBR mix. Three technical replicates were used to generate a graph and the experiments were independently performed three times with similar results. Primers used for qRT-PCR are listed in Supplementary Table 2.

### Statistical analysis and graphics

Statistical analyses were performed using R software (R v4.0.2 and RStudio v1.4.1106) or built-in statistic analyzer in GraphPad Prism software (v9.4.1). All of box-whisker plots and bar graphs were generated by GraphPad Prism. Time series plots were created using ggplot2 and the geom_ribbon function of the tidyverse library. Images were assembled in Inkscape (v1.0). All experiments included sufficient biological replicates and repeated at least twice with a similar result.

Structure prediction of AFB1 protein (Q9ZR12) was obtained from AlphaFold DB^21,22^and visualized using MolStar viewer (https://molstar.org/viewer/).

