## Supplemental Movie Legends for "The AFB1 auxin receptor controls the cytoplasmic auxin response pathway in *Arabidopsis thaliana*"

**Supplementary movie 1**: Gravitropic response of Col-0, *afb1-3*, *tir1afb2*, and *tir1afb345* roots. Time in minutes after gravistimulation is indicated.

**Supplementary movie 2:** Auxin dependent calcium transients in Col-0 and *afb1-3* roots. The decrease in background signal indicates the timepoint of 150 nM IAA treatment. Calcium transients visualized by R-GECO1.

**Supplementary movie 3**: cvxIAA triggers calcium transients in *ccvAFB1-mVenus* line, but not in *ccvTIR1* or the control *R-GECO1* lines. The decrease in background signal indicates the timepoint of 500 nM cvxIAA treatment. Calcium transients visualized by R-GECO1.

**Supplementary movie 4**: Auxin dependent calcium transients visualized by R-GECO1 in *tir1afb2* after auxin (150 nM IAA) application. The decrease in background signal indicates the timepoint of IAA treatment.

**Supplementary movie 5**: *iATTT-mCitrine*, but not *fbATTT-mScarlet*, is able to elicit auxin (150 nM IAA) dependent calcium transients. The decrease in background signal indicates the timepoint of 150 nM IAA treatment. Calcium transients visualized by R-GECO1.

**Supplementary movie 6**: *iATTT-mCitrine*, but not *fbATTT-mScarlet*, is able to rescue the gravitropic phenotype of the *afb1-3* mutant. Time in minutes after gravistimulation is indicated.
