## Supplemental Table 1 for "The AFB1 auxin receptor controls the cytoplasmic auxin response pathway in *Arabidopsis thaliana*"

**Supplementary Table 1**

| **Line** | **promoter** | **CDS** | **Tag** | **background** | **Used in figure** |
| --- | --- | --- | --- | --- | --- |
| *gAFB1-mCitrine*  *(AAAA)* | *AFB1* | *AFB1* | *mCitrine* | *afb1-3* | Fig.3b, Fig.4b,c Fig.5a,b,c,d  Supplementary Fig.1c,d,e |
| *gAFB1-NLS-mCitrine* | *AFB1* | *AFB1* | *mCitrine* | *afb1-3* | Fig.3a,b, Fig.5a,b,c,d  Supplementary Fig.3b |
| *gAFB1-NES-mCitrine* | *AFB1* | *AFB1* | *mCitrine* | *afb1-3* | Fig.3a,b, Fig.5a,b,c,d  Supplementary Fig.3b |
| *TAAA,ATAA,AATA,AAAT* | *AFB1* | *AFB1/TIR1* | *mCitrine* | *afb1-3* | Fig.4b,c |
| *iATTT* | *AFB1* | *AFB1/TIR1* | *mCitrine* | *afb1-3* | Fig.4d,e,h |
| *fbATTT* | *TIR1* | *AFB1/TIR1* | *mScarlet* | *afb1-3* | Fig.4d,e,h |
| *TTTT*  *(pAFB1:TIR1-mCitrine)* | *AFB1* | *TIR1* | *mCitrine* | *afb1-3* | Fig.4b,c  Supplementary Fig.1a,b |
| *pTIR1:TIR1-mCitrine* | *TIR1* | *TIR1* | *mCitrine* | *tir1-1* | Supplementary Fig.1a |
| *gAFB1-mCitrine cul1-6* | *AFB1* | *AFB1* | *mCitrine* | *cul1-6* | Fig.3g |
| *ccvAFB1-mScarlet* | *TIR1* | *ccvAFB1* | *mScarlet* | *afb1-3* | Fig.1f,g,h |
| *ccvTIR1-mScarlet* | *TIR1* | *ccvTIR1* | *mScarlet* | Col-0 | Fig.3d, Supplementary Fig.2c |
| *ccvTIR1 E12K-mScarlet* | *TIR1* | *ccvTIR1* | *mScarlet* | Col-0 | Fig.3c,d, Supplementary Fig.2b |
| *R-GECO1* | *UBQ10* | *R-GECO1* |  | *tir1afb2* | Fig.2d |
